# Small-scale soil microbial community heterogeneity linked to landforms on King George Island, maritime Antarctica

**DOI:** 10.1101/310490

**Authors:** Yumin Zhang, Lu Lu, Xulu xChang, Fan Jiang, Xiangdong Gao, Fang Peng

## Abstract

We analysed soil-borne microbial (bacterial, archaeal, and fungal) communities around the Fildes Region of King George Island, maritime Antarctica, which were divided into two groups according to soil elemental compositions and environmental attributes (soil chemical parameters and vegetation conditions) located in Holocene raised beach and Tertiary volcanic stratigraphy. Prokaryotic communities of the two groups were well separated; they predominantly correlated with soil elemental compositions, and were secondly correlated with environmental attributes (e.g., soil pH, total organic carbon, 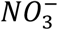, and vegetation coverage; Pearson test, *r* = 0.59 vs. 0.52, both *P* < 0.01). The relatively high abundance of P, S, Cl, and Br in Group 1 was likely due to landform uplift. Lithophile-elements (Si, Al, Ca, Sr, Ti, V, and Fe) correlated with prokaryotic communities in Group 2 may originate from weathering of Tertiary volcanic rock. The elements and nutrients accumulated during formation of different landforms influenced the development of soils, plant growth, and microbial communities, and resulted in small-scale spatially heterogeneous biological distributions. We propose that the geological evolution of the Fildes Region was crucial to its microbial community development.

**IMPORTANCE:** This current study analyzed soil-borne microbial communities around the Fildes Region of King George Island, maritime Antarctica, which were divided into two groups according to soil elemental compositions and environmental attributes. We provide new evidence for the crucial influence of landforms on small-scale structures and spatial heterogeneity of soil microbial communities.

## Introduction

Investigating microbial communities at different spatial scales, and the factors that affect microorganism distributions, are fundamental aspects of microbial biogeography (39, 42). In many terrestrial ecosystems, bacterial, fungal, and archaeal communities are distributed along soil parameter gradients (e.g., temperature, pH, water content, salinity, and nutrition; (35, 37, 48, 60, 71). Meanwhile, plants and animals that depend on the soil ecosystem may also have significant influences on microorganisms (22, 55, 75). It is therefore difficult to determine the most sensitive factors influencing microbial communities. Nonetheless, in most distinct terrestrial areas with special environments, distribution trends in the microbial community are dominantly shaped by environmental factors that limit or prevent cell growth (33, 39). Historical contingencies are also important for microbial community distribution on spatial scales of one to tens of thousands of kilometres, but such effects may be overwhelmed at small and intermediate scales (100m-1000km), including across Antarctica (12, 42). Therefore, microbial spatial structures can be used to partly reflect the external or intrinsic drivers of microbial population development and activities (72).

The extreme conditions of Antarctica, such as low temperatures, low nutrient availability, high UV radiation, and frequent freeze-thaw (14, 65), result in relatively simple ecosystems. Hence, the relatively uncomplicated food-web structure of Antarctic terrestrial habitats provides an appropriately manageable system to investigate the drivers of soil microbial diversity and composition (70). Unsurprisingly, spatial microbial community patterns have been observed here, ranging from site-specific regions to large regional scale (11, 63, 69). However, to the best of our knowledge the effects of historical contingency on microbial community distribution at small scale spatial have not been observed yet.

In this study, we used Illumina Miseq pyrosequencing and the phospholipid fatty acids (PLFA) method to survey the diversity and structure of prokaryotic and fungal communities in 12 quadrat plots around the Fildes Region, King George Island. Fildes is one of the largest ice-free regions in maritime Antarctica, and has higher biodiversity than continental Antarctica. This typical small-scale spatial region includes two Antarctic Special Protected Areas (ASPAs), covering approximately 30 km^2^ of the Fildes Peninsula, Ardley Island, and adjacent islands (8). Nevertheless, after the last glacial maximum, this region experienced multiple geologic and glacial events, including deglaciation (8400~5500 BP), glacial re-advance (after 6000 BP),

Holocene glacio-isostatic and tectonic uplift during the glacial erosive phase, and glacier retraction (30, 44). Glacial activities and past sea level changes were the key drivers of landform and soil development across the Fildes region (5, 62), and their effects on terrestrial microbial communities should not be ignored (59).

The 12 permanent quadrats analysed in our study have been established since 2013. Their primary aim was to evaluate long-term ecosystem evolution according to biomass and diversity under climate change conditions, and to build a comprehensive research platform for multi-disciplinary study, including botany, microbiology, ecology, and environmental science (68). For these purposes, all selected quadrats must: (a) include Antarctic hairgrass (*Deschampsia antarctica*), the only advanced plant discovered in the Fildes region, associated with moss and lichen; (b) have stable soil and vegetation for long term monitoring; and (c) be protected from human disturbance (mostly scientific explorers) and animal activity as much as possible. Soil maturation and vegetation colonization takes a very long time under unfavourable conditions, so the established quadrats represented natural and stable habitats around the region. Recently, based on 454 pyrosequencing data, Wang et al.(61) found that the diversity and structure of soil bacterial communities in four sites of the Fildes region were significantly affected by pH, phosphate phosphorus, organic carbon, and organic nitrogen. The relationships between microbial communities, geological factors, and landform development have not been studied.

In this study, we attempt to answer three questions: (i) what is the microbial community structure in this small-scale region of maritime Antarctica; (ii) which factors affect the distribution of microbial communities; and (iii) do microbial communities reflect different landform development in history. In order to explore these issues, a number of soil chemical properties and vegetation attributes were measured, representing conventional environmental factors; the soil element composition was determined by X-ray fluorescence spectrometer, which is an acceptable proxy for soil or sediment erosion and development in different landforms (4, 28, 66), and combining geological literature related to the Fildes region to represent landform types. We aim to improve understanding of terrestrial microbial communities in maritime Antarctic ice-free areas, and contribute to a new perspective on small-scale microbial biogeography.

## Results

### Soil elemental compositions and environmental attributes of quadrats

A total of 20 elements in the sample soils were detected by X-Ray fluorescence spectrometer, and 11 environmental attributes were measured (Table S1). Principle component analysis with normalized whole soil elemental compositions and environmental attribute data showed that 36 samples were well separated by the original point of the PC1 coordinate axis (Fig. 1a). Hence the quadrat plots could be divided into two groups reflecting different soil types and environmental conditions, in which Group 1 included quadrat plots Q2, Q3, Q6, and Q7, and Group 2 included all other quadrat plots (Q1, Q4, Q5, Q9, Q10, Q11, Q12, and Q13). The heatmap cluster analysis supported the grouping suggested by whole element and environmental data (Fig. 1b).

**Fig. 1.**
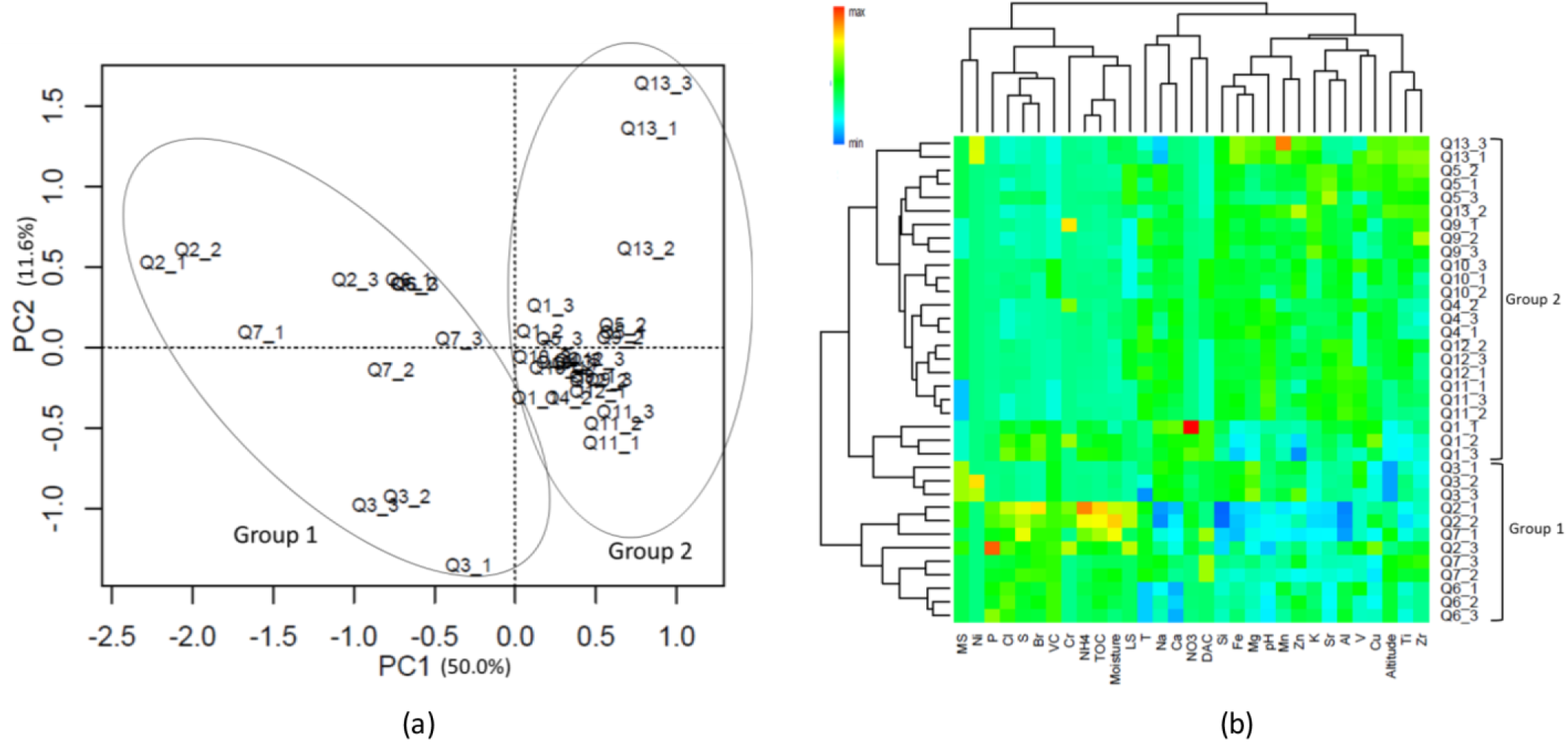
(a) Principle component analysis (PCA) and (b) heatmap cluster analysis of the normalized soil elemental compositions and environmental attribute data. The values of PC1 and 2 are percentages of total variations that can be attributed to the corresponding axis. Abbreviations: T, temperature; TOC, total organic carbon; MS, moss species amount; LS, lichen species amount; DAC, hairgrass (*Deschampsia antarctica*) coverage; and VC, total vegetation coverage.

The soil element profile revealed that lithophile elements (Si, Al, Ca, Mg, Fe) occupied the major portions. Twelve elements (Al, Ca, Cu, Fe, K, Mg, Mn, Si, Sr, Ti, V, and Zn) were more abundant in Group 2, and four elements (P, S, Cl, and Br) were significantly more abundant in Group 1 (Wilcoxon test, *P* < 0.05, Table S1). PERMANOVA analysis revealed a highly significant difference in soil element composition between the two sample groups (*Pseudo-F* = 17.74, *P* < 0.01). Pairwise correlative comparisons between elements demonstrated that P, S, Cl, and Br were positively correlated with each other, and negatively correlated with Mg, Al, Si, Ca, Mn, Zn, Sr, V, Fe and Ti, which also had positive correlations with each other (Fig. S1). A significant difference was also observed in the environmental attributes between Group 1 and Group 2 (PERMANOVA test, *Pseudo-F* = 15.17, *P* < 0.01), consisting of lower soil pH and higher total organic carbon (*TOC*, 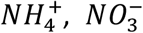) and moisture contents in Group 1. In addition, vegetation properties such as hairgrass coverage (DAC) and total vegetation coverage (VC) were also higher in Group 1 plots (Wilcoxon test, *P* < 0.05, Table S1).

### Diversity and composition of microbial communities in quadrats

After sequence-quality filtering, we obtained a total of 2,389,662 high-quality bacterial 16S rRNA gene reads, 1,423,619 archaea 16S rRNA gene reads, and 1,953,908 ITS reads. These reads constituted 98,887, 49,000, and 8,464 operational taxonomic units (OTUs) at a 0.03 discrepancy (97% identity) for bacterial, archaeal, and fungal taxa, respectively. The OTUs diversities of Shannon, Chao 1, and ACE index of bacteria, archaea, and fungi did not differ between Group 1 and Group 2 (Wilcoxon test, *P* > 0.05, Table S2). For bacteria, 20 phyla and some unidentified bacteria (0~0.8%) were detected, and the OTU sequences of most quadrat soils were dominated by *Actinobacteria* (24.2%), *Acidobacteria* (14.7%), *Proteobacteria* (15.1%), *Chloroflexi* (12.3%), and *Gemmatimonadetes* (7.2%) (Fig. S2). For archaea, a number of OTU sequences (74.6%, maximum) were not assigned to any taxon; the remainder was dominated by *Crenarchaeota* (94.5%) and *Euryarchaeota* (5.5%), and the dominant classes of the phyla were *Thaumarchaeota* and *Thermoplasmata*, respectively. For fungi, six phyla were detected, and all quadrat plot soils were dominated by *Ascomycota* (69.1%), *Basidiomycota* (17.6%), and *Zygomycota* (4.5%). The percentage of unassigned OTUs and unidentified fungi were 6.8%–64.3% and 0.4%–7.7%, respectively. PERMANOVA tests of Bray-Curtis distances revealed significant differences between the two groups for prokaryotic 16S rRNA genes (integrated data normalized by the bacterial and archaeal OTU data set, Pseudo-F = 3.1, *P* = 0.0002), but not for fungal ITS genes (Pseudo-F = 1.3, *P* = 0.196). This was consistent with the grouping result of nonmetric multidimensional scaling (NMDS) analysis with the bacterial and archaeal OTU dataset (Fig. 2).

**Fig. 2.**
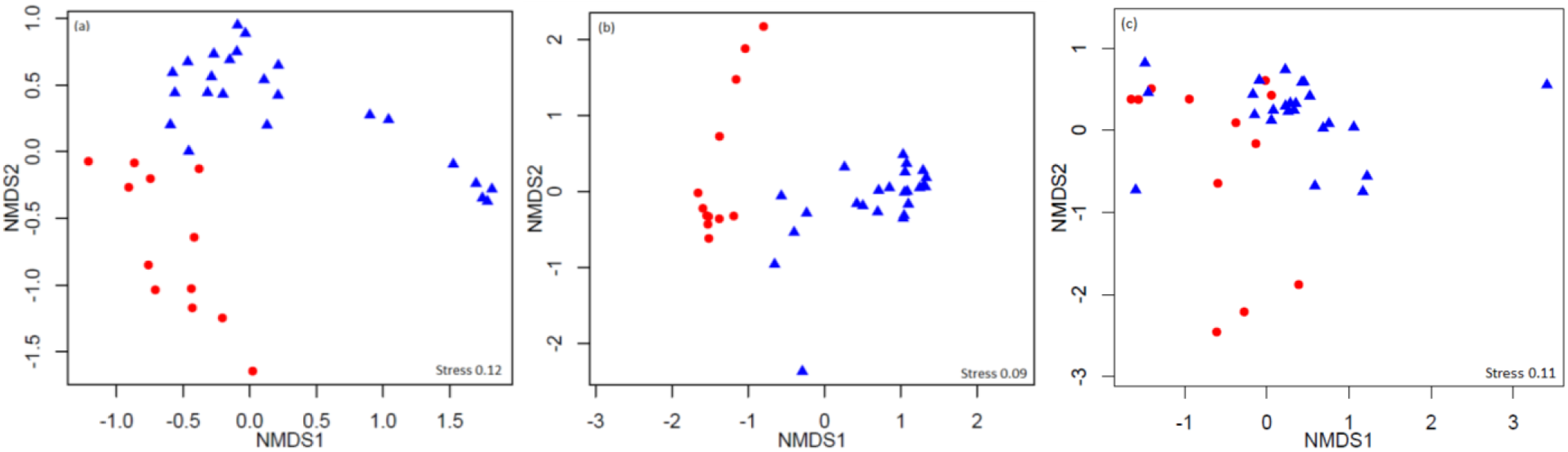
Nonmetric multidimensional scaling (NMDS) analysis of the (a) bacterial, (b) archaeal, and (c) fungal Operational Taxonomic Unit (OTU) datasets. Circle = Group 1; triangle = Group 2.

### Links among microbial composition, soil elemental composition, and environmental attributes

The prokaryotic OTU composition showed a strong and significant correlation with soil elements (r = 0.59, *P* < 0.01, Pearson test), and a less but still significant correlation with environmental attributes (r = 0.52, *P* < 0.01, Pearson test; Fig. 3a). Fungal community composition did not show strong correlations with either soil elements or environmental attributes (P > 0.05; Fig. 3b). The soil elements and environmental attributes in canonical correspondence analysis (CCA) were selected by variation inflation test (see the Methods for details). CCA results for bacterial and archaeal community composition and soil elements, with significant models at the confidence level (both *P* < 0.01), indicated that the 11 soil elements are important factors controlling bacterial and archaeal community structures, and explain 60.0% and 47.3% of their variations, respectively (Fig. 4a and b).

**Fig. 3.**
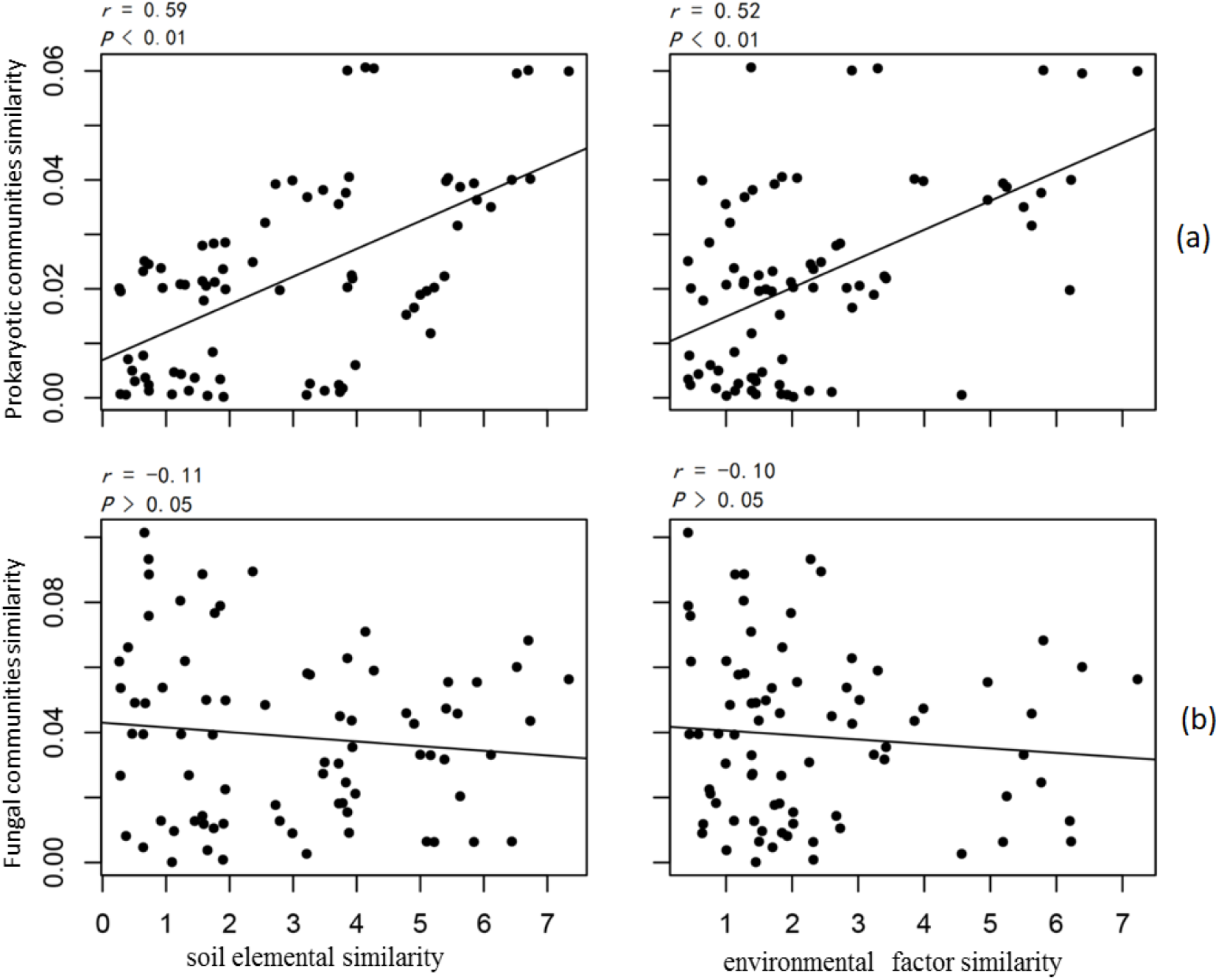
Pearson correlations between (a) the prokaryotic community and (b) the fungal community with soil elemental compositions and environmental attributes. Similarity values are directly indicated by calculated pairwise Euclid distances between samples.

**Fig. 4.**
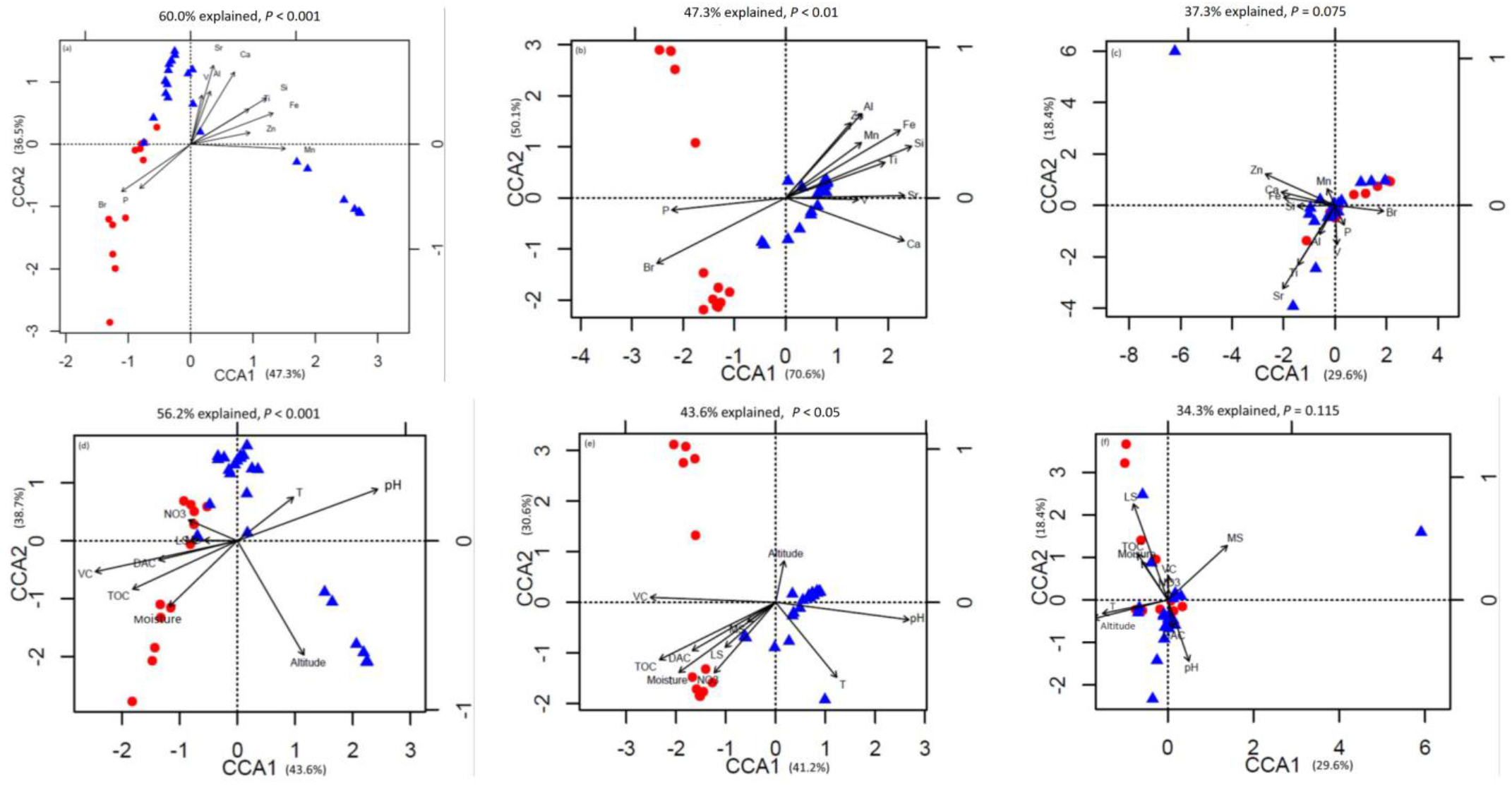
Canonical correspondence analysis (CCA) of (a) Bacterial Operational Taxonomic Unit (OTU) data and elemental compositions; (b) archaeal OTU data and elemental compositions; (c) fungal OTU data and elemental compositions; (d) bacterial OTU data and environmental attributes; (e) archaeal OTU data and environmental attributes; (f) fungal OTU data and environmental attributes.

Among these elements, P and Br were important elements controlling microbial community structures in Group 1, and other lithophile and metal elements controlled Group 2. The importance of these soil elements was verified by a Monte Carlo test (P < 0.05, 999 permutations) with prokaryotic community data comprising bacterial and archaeal community compositions (Table 1). For fungal communities, CCA analysis showed that the two groups were not well separated from the others, and the model was not significant within the confidence level (P > 0.05). Only 37.3% of fungal community variations could be explained by the 11 soil elements (Fig. 4c). Considering the relationships between microbial communities and environmental attributes, the same analysis was made by CCA (Fig. 4d, e, and f). The models were significant between both bacterial (P < 0.01, 56.2% explained) and archaeal (P < 0.05, 43.6% explained) community structures and environmental attributes. The Monte Carlo test (999) revealed that total organic C (TOC), soil pH, moisture, site altitude, hairgrass coverage (DAC), and total vegetation coverage (VC) showed strong effects on prokaryotic communities. For fungal communities, the model was not significant within the confidence level (P > 0.05, 34.3% explained); however, soil pH, site altitude, moss species amount (MS), and lichen species amount (LS) were the environmental attributes affecting fungal community composition (Monte Carlo test, *P* < 0.05, 999 permutations; Table 1). For the mantel test of microbial community structures, including all factors investigated in this study, see Table S3.

**Table 1.** Monte Carlo test of the factors (soil elemental compositions and environmental attributes) and compositions of microbial communities and phospholipid fatty acids (PLFA). Significant differences (P < 0.05) are indicated in bold. ***P < 0.001, **P < 0.01, *P < 0.05. P-values based on 999 permutations.

| Soil elements | Prokaryote |  | Fungi |  | PLFA |  |
| --- | --- | --- | --- | --- | --- | --- |
|  | r2 | *P*-value | r2 | *P*-value | r2 | *P*-value |
| Al | 0.2547 | 0.002** | 0.0014 | 0.979 | 0.4648 | 0.001*** |
| Ca | 0.5647 | 0.001*** | 0.0012 | 0.981 | 0.3276 | 0.003** |
| Mn | 0.5235 | 0.001*** | 0.0018 | 0.965 | 0.3859 | 0.001*** |
| P | 0.3372 | 0.005** | 0.0488 | 0.378 | 0.2883 | 0.012* |
| Si | 0.5621 | 0.001*** | 0.0299 | 0.609 | 0.6624 | 0.001*** |
| Sr | 0.5332 | 0.001*** | 0.2784 | 0.007* | 0.4546 | 0.001*** |
| Ti | 0.3316 | 0.003** | 0.2702 | 0.010* | 0.3503 | 0.001*** |
| V | 0.2005 | 0.020* | 0.0408 | 0.697 | 0.1828 | 0.041* |
| Zn | 0.2309 | 0.009** | 0.1120 | 0.216 | 0.3289 | 0.003** |
| Br | 0.5059 | 0.001*** | 0.0369 | 0.558 | 0.6223 | 0.001*** |
| Fe | 0.5210 | 0.001*** | 0.0165 | 0.761 | 0.6978 | 0.001*** |
| Environmental factors | Prokaryote |  | Fungi |  | PLFA |  |
|  | r2 | *P*-value | r2 | *P*-value | r2 | *P*-value |
| TOC | 0.4139 | 0.001*** | 0.1748 | 0.099 | 0.6776 | 0.001*** |
| NO3 | 0.0667 | 0.286 | 0.0055 | 0.792 | 0.5661 | 0.120 |
| T | 0.1531 | 0.068 | 0.1867 | 0.055 | 0.1042 | 0.176 |
| pH | 0.6958 | 0.001*** | 0.2385 | 0.007** | 0.4830 | 0.001*** |
| Moisture | 0.2644 | 0.012* | 0.1231 | 0.128 | 0.5844 | 0.001*** |
| Altitude | 0.3003 | 0.002** | 0.2463 | 0.024* | 0.0065 | 0.912 |
| DAC | 0.2007 | 0.030* | 0.0454 | 0.516 | 0.3487 | 0.003** |
| MS | 0.0320 | 0.557 | 0.2723 | 0.007** | 0.0505 | 0.430 |
| LS | 0.0624 | 0.325 | 0.5927 | 0.001*** | 0.1367 | 0.103 |
| VC | 0.6465 | 0.001*** | 0.0298 | 0.779 | 0.4422 | 0.001*** |

### Microbial biomass and microbial diversity determined by the phospholipid fatty acids (PLFA) method

The total amounts of PLFA (totPLFA) of Group 1 were significantly higher than those of Group 2 (Wilcoxon test, *P* < 0.05; Fig. S3). CCA analysis of the individual relative concentration (mol%) of the 45 most common PLFAs showed that, on the whole, the 11 soil elements and the 11 environmental attributes were all important factors controlling soil PLFA patterns (Fig. S4, *P* < 0.01), with 47.5% and 47.0% of the variations explained, respectively. Among these factors, each of the 11 elements, and pH, moisture, total organic C (TOC), *Deschampsia antarctica* coverage (DOC), and total vegetation coverage (VC) of the environmental attributes had significant effects on soil PLFA composition (Table 1). Microorganism categories including bacteria, fungi and protozoa were classified by indictor PLFAs according to microbial identification systems (MIDI). The relative abundance of AM fungi, actinomycetes, and anaerobes were higher in Group 1, and Gram-negative bacteria was higher in Group 2 (Fig. S5; Welch’s t-test, two-sided, *P* < 0.05).

### Differences of microbial community composition between the two groups

In our analysis, the classified mode of the Random forests machine learning technique (9, 16) could be accepted if the ratio of the baseline error to the observed error was greater than 2, and we considered an OTU to be highly predictive if its importance score was at least 0.001. For bacteria, random forest analysis revealed that 58 OTUs distinguished the two groups, *Acidobacteria* were overrepresented in Group 1, and the OTUs assigned to the *Thermoleophilia* class of the phylum *Actinobacteria*, and genus *Geobacillus* of phylum *Firmicutes*, were overrepresented in Group 2. For archaea, 38 OTUs distinguished the two groups, except for 12 OTUs with no assigned taxa. Some 11 OTUs were overrepresented in Group 1 and 17 OTUs were overrepresented in Group 2; all were assigned to genus *Candidatus Nitrososphaera* of phylum *Crenarchaeota.* As the ratio of the baseline error to the observed error of the random forest analysis with fungal OTUs was less than 2, we considered that the non-obvious classified result suggested that there was no credible difference in fungal community composition between the two groups (Table S4).

## Discussion

The prokaryotic community composition of the quadrats can be divided into two groups that correlate with soil element compositions and environmental attributes. Interestingly, a published geologic map of the Fildes region and previously reported literature (38, 44) showed that the quadrats in Group 2 were located in Tertiary volcanic stratigraphy, and those in Group 1 were found on a Holocene raised beach (Fig. 5). We suggest that there is a potential relationship between microbial communities and the development of landforms at this small spatial scale.

**Fig. 5.**
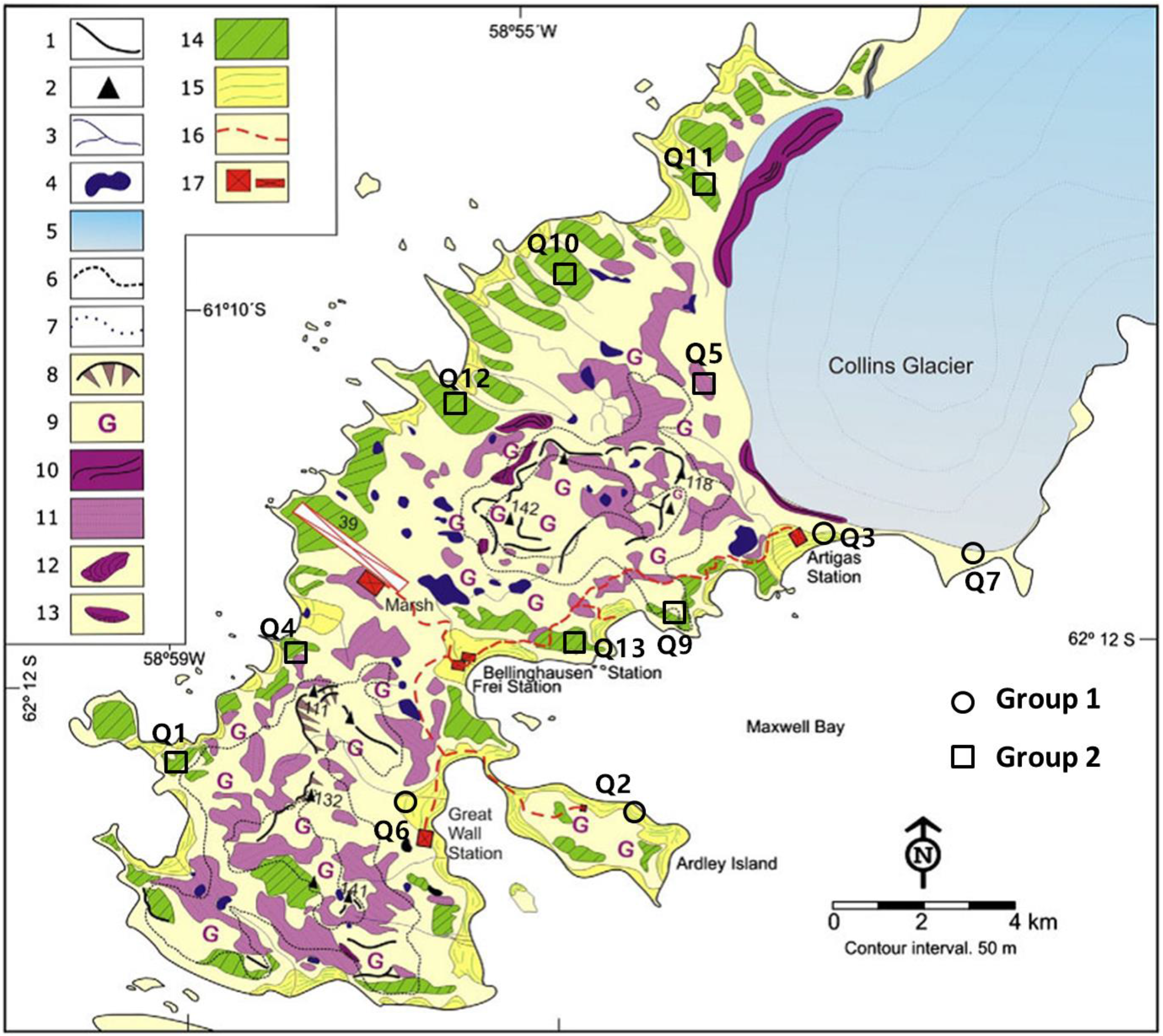
Geomorphological map of the Fildes region, derived from Michel et al (44). Quadrats of Group 1 and Group 2 were located at Holocene raised beaches (No. 15) and marine platforms (periglacial landforms belonging to Tertiary volcanic stratigraphy, No. 14). The landform type of quadrat Q7 can be deemed as part of the Holocene raised beaches because it suffered recent glacio-isostatic uplift but was still covered by ice during that uplift (personal communication; Michel, 2018). Please refer to original literature for landforms marked by other numbers.

In this area, a series of geological events, including volcanic activity, glacial erosion and retraction, isostatic uplift, and sea level change, created rich landform types. According to geomorphological and sedimentary evidence, relative sea level (RSL) gradually fell to < 14.5 m between 7000 and 4750 cal a BP as a consequence of isostatic uplift in response to regional deglaciation (30, 62). During landform formation, rich marine elements and nutrients were transferred to the land (5, 44). Moreover, from approximately 2500 years ago, mammals, especially penguins, began to colonize the newly uplifted beaches until at least ~500 years ago when the raised beaches were abandoned (according to chronological research of abandoned rookeries on King George Island (54)). These abandoned penguin rookeries are indicators of Holocene paleoclimate and also accumulated rich nutrients during the period (3). These input elements, nutrients, and marine microorganisms clearly promoted the development of soil and plant growth, and influenced patterns of microbial community formation.

Soil elemental profiles can be seen as proxy indicators of soil types and landforms with strong soil-landform relationships (20, 34). In previous studies, soil compositions tested by X-ray fluorescence spectrometer have been revealed to be the key factors for the distribution of bacterial and fungal communities in some field sites, sediments, glacier forefields, and deserts (27, 29, 43, 52). In this study, the CCA analysis and similarity test showed that both environmental attributes and soil element compositions could influence the microbial structure and biomass. However, compared with environmental attributes, the relationship between soil element composition and prokaryotic community was stronger (Fig 4; Fig 5). Mantel analysis revealed that the relative abundance of almost every element was important for shaping prokaryotic compositions (Table S3). The quadrat plots located on the Holocene raised beach landform showed relatively high abundances of P, S, Cl, and Br, which were more correlated to marine environments and organisms. These elements are readily absorbed by vegetation and microorganisms, and presumably resulted in the development of microbial community structures in Group 1 (CCA analysis; Fig. 4). The accumulation of elements P and S may represent marine input but also mammal and bird excrement that accumulated in these raised beaches during the early stage of uplifted landform formation (54). Meanwhile, the halogen elements Cl and Br in island coastal soil likely derived mostly from seawater (49).

Conversely, the bacterial and archaeal community compositions of Group 2 were more correlated with lithophile-elements (Si, Al, Ca, Sr, Ti, V, and Fe, CCA analysis), and the landforms were almost completely isolated from the external environment until the icecap retreated ~11000-7500 cal a BP (62). This suggests that soil in quadrats located in tertiary volcanic stratigraphy mainly developed from the chemical and biological weathering of volcanic rock generated by Tertiary volcanism, and underwent paraglacial and periglacial processes. That may explain the lower soil biomass, nutrition, and vegetation coverage, as compared with Group 1; the limited nutrient input distinguished the prokaryote community composition from that of the nutrient-rich soil of Group 1. Therefore, we believe that the element composition of the soil associated with these landforms reveals geological background and historic effects.

In Group 1, the soil contained high contents of TOC, 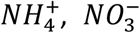) and vegetation coverage, which also correlates with prokaryotic community. The relatively low pH values (Table S1) may be a result of higher vegetation coverage with more humus and fulvic acids produced by moss and lichen (21). As the rich nutrients and elements transferred from Holocene raised beach marine environments could promote soil development and plant growth, these environmental attributes seem to be a secondary factor affecting the prokaryotic community when compared to soil element compositions. Unlike Group 1, very small amounts of nutrients in the soil samples of Group 2 were more likely caused by current precipitation, snowfall, and animal activity. In keeping with reported studies (19, 23, 35), pH is one of the most influential factors affecting the distribution of microbial communities in this study.

Interestingly, both prokaryotes and fungi communities were significantly correlated to the altitude of the sample location. Despite the slightly different altitudes (ranging from 11-56 m), they do not lead to significant changes in temperature, oxygen content, etc., which seems to suggest that geological uplift had an impact on microbial communities. In addition, moss and lichen species were significantly associated with fungal communities. It was previously reported that some fungal species coexist with moss and lichen in Antarctica (36, 58). We also noted that the soil element compositions and environmental attributes of ancient landforms investigated in our study were relatively stable, while those of younger landforms were more volatile (from Euclidean distances computed between samples from the PCA analysis in Fig 1a, Table S5). This suggests that the quadrat plots of Group 1 may be in an unstable new geological layer within a transboundary ecological stage from ocean to land, and disturbance from the new terrestrial environment may increase the heterogeneity of the geomorphic ecology.

As the quadrats in our study all had hairgrass growth, vegetation may be one of the main causes of this difference. Thus, our results were similar to bacterial community compositions in other vegetated parts of Antarctica (36, 58), with relatively high abundances of *Chloroflexi* and *Gemmatimonadetes*, which have strong reported relationships with plants (2, 10, 17). Previous studies of Antarctic archaeal communities were mostly concentrated in marine and lake environments (13, 18, 31, 32, 45), and in this study, only two archaeal phyla (*Crenarchaeota* and *Euryarchaeota*) were detected, with *Crenarchaeota* representing the overwhelming majority (> 90%) of archaeal communities. This was consistent with other terrestrial archaeal structures of Antarctica derived using other investigated methods (e.g., clone libraries of rRNA genes and microarray (1, 70).

Random forest analysis revealed that OTUs belonging to *Alphaproteobacteria, Acidobacteria*, and *Bacteroidetes* were mostly overrepresented in Group 1. These phyla have shown positive correlations with vegetation and the rhizosphere in farmland, arctic glacier moraines, and the Brazilian Antarctic Station (41, 55, 57). The results of our study showed different patterns at the family level, with *Acidobacteriaceae, Koribacteraceae, Chitinophagaceae*, and *Rhodospirillaceae* the most overrepresented families in Group 1. The family level differences from our study could be due to the locations of sampling points and the diverse sequencing methods. Conversely, in Group 2, the major overrepresented OTUs were class *Thermoleophilia* of phylum *Actinobacteria. Thermoleophilia* is a newly proposed class of phylum *Actinobacteria* that was created from the splitting of *Rubrobacteridae* (40), and its ecological position is not well understood. However, *Thermoleophilia* is abundant in deserts and glacier forelands (15, 74), and some isolated parts could be cultured in low nutritional media during long incubation periods. Thus, it is reasonable that this class is found in the quadrats located in volcanic stratigraphy with high proportions of lithospheric elements and low nutrition conditions. In addition, we also found that five OTU sequences affiliated to *Flavobacteriaceae* extracted from Group 1 were clustered in marine clades, and no marine clade OTUs of *Flavobacteriaceae* were found in Group 2 (Fig. S6). Members of the family *Flavobacteriaceae* are among the most abundant picoplankton in coastal and polar oceans, and a number of genera have potential evolutionary sources from the ocean (7). Regarding genus *Candidatus Nitrososphaera*, the vital ammonia-oxidizing archaea (51, 53) was the overrepresented archaeal OTU in both groups. The uncultivable species *s_Ca.* N. SCA1170 was a major genus in Group 2 but did not appear in Group 1. This, along with evidence from NMDS analysis, implies that the two different landforms have diverse archaeal communities. Those suggested that spatial constraints for microorganisms also occur at small spatial scales.

The importance of geological factors, such as the landform and lithology, on microbial structure is less well understood (59). Locations with distinct geologic factors generally exhibit geographical isolation; hence, they are mostly distributed at large and global scales. Limited research has shown that different landforms and soil profiles are also important drivers of bacterial diversity at the regional scale (>1000 km distance), and their impacts are more significant than contemporary environmental factors (25, 47). Interestingly, we found that, on such a small spatial scale, prokaryotic communities also showed a landform-governed distribution trend, and the microbial community structure is expected to be an indicator of the formation of the landform. The role of geological evolution in microbial distribution can be highlighted in this study because: (i) a clear effect of the geological evolution of the Fildes region in maritime Antarctica. Glacial activity, sea level changes, and tectonic uplift due to climate change after LGM have all resulted in landform heterogeneity at a small spatial scale; (ii) seasonal freezing-thawing cycles in the area have enhanced soil development, and promoted soil particle and nutrition migration to upper and surface soil layers; and (iii) the low activity of microorganisms under the cold climate, and less human disturbance where the quadrats were established, maintained relatively stable microbial community diversity for a long time after the geological changes. However, in contrast to other research, we did not attempt to classify the common environmental factors measured in this study as ‘contemporary environmental’ factors because those representing soil nutritional conditions were considered to be the consequence of landform development (44, 54), especially in Group 1. In our study, the environmental factors had strong influences on bacterial and archaeal community structures. Nonetheless, these were likely to play subsequently important roles in the distribution of microbial communities, predominantly driven by landforms and soil element compositions.

In conclusion, this study provides evidence for the influence of geological evolution on the small-scale distribution of microbial communities. As a result, microbial community structure is proposed as an indicator of the two different landforms in the Fildes region, King George Island. In addition, other locations in Antarctica experience the same type of glacial activity and isostatic uplift as the coastal ice-free areas around King George Island, maritime Antarctica, and Prince Charles Mountains area, East Antarctica (64), implying that microbial communities may also be diverse and influenced by different geological evolution events at small to moderate spatial scales in these areas. Continued research already, in progress, will verify whether microbial communities can be used as indicators of different landforms in other, similar geological areas in maritime Antarctica. This will contribute to finding different microbial communities in limited spatial regions based on geological research, and will examine different types of geological heterogeneity according to microbial communities.

## Materials and Methods

### Quadrat plot description, soil sampling, and sample preparation

The Fildes region is the largest ice-free area on King George Island, with a humid and relatively mild sub-Antarctic maritime climate. The mean annual temperature and precipitation are −2.4°C and over 500 mm, respectively (26). The 12 permanent quadrat plots (1.5 m × 1.0 m each) investigated in this study had been established on Fildes Peninsula and Ardley Island between 2013 and 2015. For plot characteristics, please see the Introduction for details. Each quadrat plot was fenced to minimize disturbance. GPS coordinates, vegetation characteristics, and the landscape of quadrat locations are shown in Table 2 and Fig. S7. The distance between quadrat plots ranges from approximately 1.6 to 8.2 km. Sampling occurred during China’s 33rd Antarctic expedition in January 2017. Soils were sampled from the A-horizon (10 cm), at an internal distance of approximately 3-5 m, in triplicate around each quadrat plot. Soil samples collected for each replicate were taken from five soil cores (5 cm diameter) and mixed thoroughly. A total of 36 soil samples were placed in sterile plastic bags, and soil DNA was extracted within 2 h in the laboratory of the Great Wall Station. The remaining soils were stored in the freezer until further soil physic-chemical property analysis.

**Table 2.**
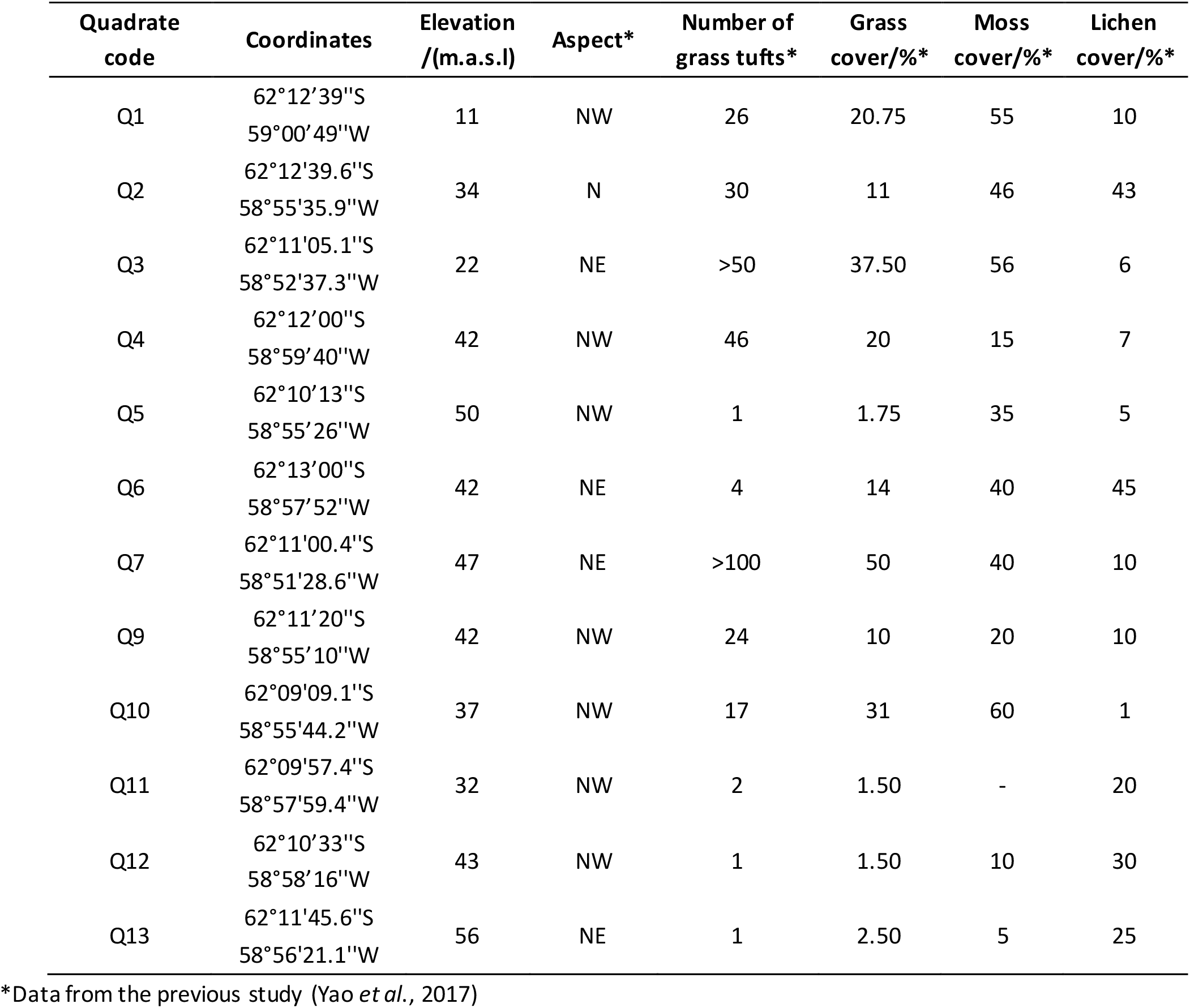
Locations and partial vegetation properties of the 12 soil quadrats.

### DNA extraction, PCR, pyrosequencing, and pyrosequencing data treatment

Genomic DNA was extracted using a PowerSoil DNA Isolation Kit (MoBio, Carlsbad, CA, USA) according to the manufacturer’s instructions. Duplicate DNA extraction was performed for each sampling plot, and all duplicated DNA products were pooled to reduce potential DNA extraction bias. Afterwards, DNA concentration was measured by UV spectrophotometer (Eppendorf, Bio Photometer), and its molecular size was estimated by 0.8% agarose gel electrophoresis. Details of pyrosequencing and pyrosequencing data treatment are described in Appendix S1. These sequence data have been submitted to the DDBJ/EMBL/GenBank databases (SRA) under accession no. SRP132288, accession no.SRP132345 and accession no.SRP132350.

### PLFA analysis

Phospholipid fatty acids (PLFAs) from soil samples were extracted, fractionated, quantified, and analysed using the protocol described by (6). In brief, 2.0 g of soil (dry weight) was extracted with a chloroform-methanol-citrate buffer mixture (1:2:0.8) and fractionated into neutral lipids, glycolipids, and phospholipids on a silicic acid column (Agilent Technologies, Sillic Box, CA, USA). Phospholipids were subjected to mild alkaline methanolysis, after separating out fatty acid methyl esters on an Agilent 6890N gas chromatograph equipped with a flame ionization detector and an HP-1 Ultra 2 capillary column (Agilent Technologies, Santa Clara, CA, USA). Peak areas were quantified by adding methyl non-adecanoate fatty acid (C19:0) (Sigma) as an internal standard. The fatty acid methyl esters were prepared according to the MIDI protocol and analysed using the MIDI Sherlock Microbial Identification System (MIDI, Newark, DE). The fatty acids i14:0, i15:0, a15:0, i16:0, a16:0, i17:0, and a17:0 represented gram-positive bacteria, 16:1ω9c, cy17:0, 18:1ω5c, 18:1ω7c, and cy19:0 represented gram-negative bacteria, 10Me16:0 (24), 10Me17:0, and 10Me18:0 represented Actinomycetea (73), branched monoenoic and mid-branched saturated fatty acid PLFAs represented anaerobic microorganisms (76), and 16:1ω5 represented AM fungi (46). PLFAs were categorized and calculated in the MIDI Sherlock Microbial Identification System (MIDI, Newark, DE).

### Soil element composition determined by X-ray fluorescence spectrometer

Soil samples were dried at 105°C for 6 h and then ground into powder. The soil powder was pressed in a 45 mm bore steel die under an approximately 20 t hydraulic press. Every soil sample formed a stable soil pie of 45 mm diameter and 10 mm height. These pies were generally analysed within a few hours. The elements within soil samples were determined by X-ray fluorescence spectrometry (Bruker AXS, Germany) using a standardless quantitative analysis method (29). We removed poor quality elemental signals that rarely appeared (< 0.01%), generally in only one or two samples.

### Soil parameters and vegetation attribute measurements

Soil temperature was measured by a plug-type thermometer (ZD Instrument, China) at depths of 15 cm during soil sampling. Soil pH was measured by adding 10 ml of distilled water to 5 g of soil, and recording pH by a pH electrode (Mettler-Toledo, Switzerland). Soil moisture was determined as the gravimetric weight loss after drying the soil at 105ºC until reaching a constant weight. Analysis of total organic carbon was performed using a TOC analyser (vario TOC, Elementar, Germany). To measure 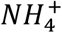 and 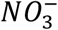, 10 g of soil was suspended in 50 ml of 2 mol/L KCl solution and shaken at 25ºC for 1 h. Then, the soil solution mixture was centrifuged for 5 min in 3000 g. Subsequently, clear supernatant was passed through a filter of 0. 45 μm (Millipore, type GP), and analysed using a continuous flowing analyser (FIAstar 5000, Foss, Denmark). Each quadrat of 1 × 1 m was selected to measure vegetation attributes including moss species number (MS), lichen species number (LS), hairgrass (*Dechampsia antarctica*) coverage (DAC), and total vegetation coverage (VC) according to previous protocols (68).

### Data statistical analyses

For estimating bacterial, archaeal, and fungal diversity, Operational Taxonomic Unit (OTU) analysis including the Shannon, Chao1, and ACE indices was performed using the Mothur v. 1.30.2 software package (50). The relationships between soil elemental compositions and environmental attributes in the 32 soil samples were analysed by principal component analysis and hierarchical clustering heatmap analysis using the R v. 3.3.1 statistical software. The Wilcoxon test was performed for the soil elements and environments to determine the level of significance with a two-sided hypothesis using the Statistical Package for the Social Sciences software (SPSS). Significant differences in soil elemental compositions, environmental attributes, and microbial community structures between groups were determined by permutational multivariate analysis of variance (PERMANOVA) on 999 permutations of residuals under a reduced model using the R v. 3.3.1 statistical software. The Bray-Curtis distance was used to obtain dissimilarity matrices in the PERMANOVA test for microbial OTU data. The similarity test, Mantel test, and Canonical correspondence analysis (CCA) were used to evaluate the linkages between microbial community structures (general levels) and soil elemental compositions and environmental attributes with the Vegan package (v. 2.4-1) in R v. 3.3.1 according to the method described by Yang et al (67). Variation inflation factors were used to select factors in CCA modelling, of which the variance of canonical coefficients was not inflated by the presence of correlations with other factors, so that soil elements and environmental attributes were removed if the variation inflation factor was larger than 20. Variation partitioning analysis resulted in 11 soil elements (Si, Ca, Zn, Fe, Al, Mn, V, Ti, Sr, P, Br) and 10 environmental attributes. The effect of factors on microbial community structures and the PLFA profile was estimated by a Monte Carlo permutation test (999 permutation). Differences in microbial categories marked by PLFA was determined using Welch’s t-test (two-sided) using the STAMP software (v. 2.1.3) package.

## Acknowledgements

This work was supported by the R & D Infrastructure and Facility Development Program of the Ministry of Science and Technology of the People’s Republic of China (Grant No. NIMR-2017-8), the National Natural Science Foundation of China (Grant No. 31270538), and the Chinese Polar Scientific Strategy Research Fund IC201706.

We also thank Roberto Michel for sharing his knowledge of the geological environment of the Fildes region.

